# Geometric effects of volume-to-surface mapping of fMRI data

**DOI:** 10.1101/2020.11.09.373860

**Authors:** Keith George Ciantar, Christine Farrugia, Kenneth Scerri, Ting Xu, Claude J. Bajada

## Abstract

In this work, we identify a problem with the process of volume-to-surface mapping of functional Magnetic Resonance Imaging (fMRI) data when interested in investigating local connectivity. We show that neighborhood correlations on the surface of the brain vary spatially with anatomical patterns (gyral structure), even when the underlying volumetric data are uncorrelated noise. This could potentially have impacted studies focusing upon local neighborhood connectivity. We explore the effects of this anomaly across varying data resolutions and surface mesh densities, and propose an approach to mitigate these unwanted effects.

## 1 Introduction

Taking a surface-based approach has become a popular choice when performing analysis of functional MRI data. Indeed, an increasing number of large projects provide surface-based data (e.g. Human Connectome Project [HCP] (Van Essen et al, 2012, 2013), Adolescent Brain Cognitive Development [ABCD] study (Bjork et al, 2017)), as well as software packages that facilitate surface analysis (e.g. FreeSurfer or Connectome Workbench). Recent fMRI packages have also implemented surface-based analysis in the preprocessing pipelines (e.g. the HCP Pipelines (Glasser et al, 2013) and fmriPrep (Esteban et al, 2019)). There are good theoretical (Brodoehl et al, 2020; Glasser et al, 2016) and practical advantages (Coalson et al, 2018) to performing fMRI analysis on a cortical surface mesh. The primary advantages are that smoothing on a two-dimensional surface respects the geometry of the brain and reduces signal contamination from cortical areas that are geodesically distant, yet whose euclidean separation is small – such as opposing banks of neighboring sulci. Additionally, cross-subject alignment is fundamentally easier using surface – rather than volumetric – registration.

As part of the surface-based preprocessing pipeline, the user must first map their volumetric (voxel) data to surface vertices. This process has the potential to introduce artefacts that may affect some downstream surface-based analysis. In this technical note, we identify and report a problem with the process of volume-to-surface mapping in the context of local neighborhood connectivity analysis. The effect has potentially impacted some studies that have not taken it into consideration.

## 2 The Problem

An anomaly was noted when analysing regional boundaries (or edges) in functional signatures across the cortex (Bajada et al, 2020). Specifically, when aiming to characterize regional boundaries by calculating functional changes between neighboring vertices along the surface, we observed a clear anatomical pattern that follows the gyral folds of the brain, even when using stochastic data as input (Figure 1). This suggests that the local connectivity analysis might be affected, at least partially, by surface geometry.

**Fig. 1.**
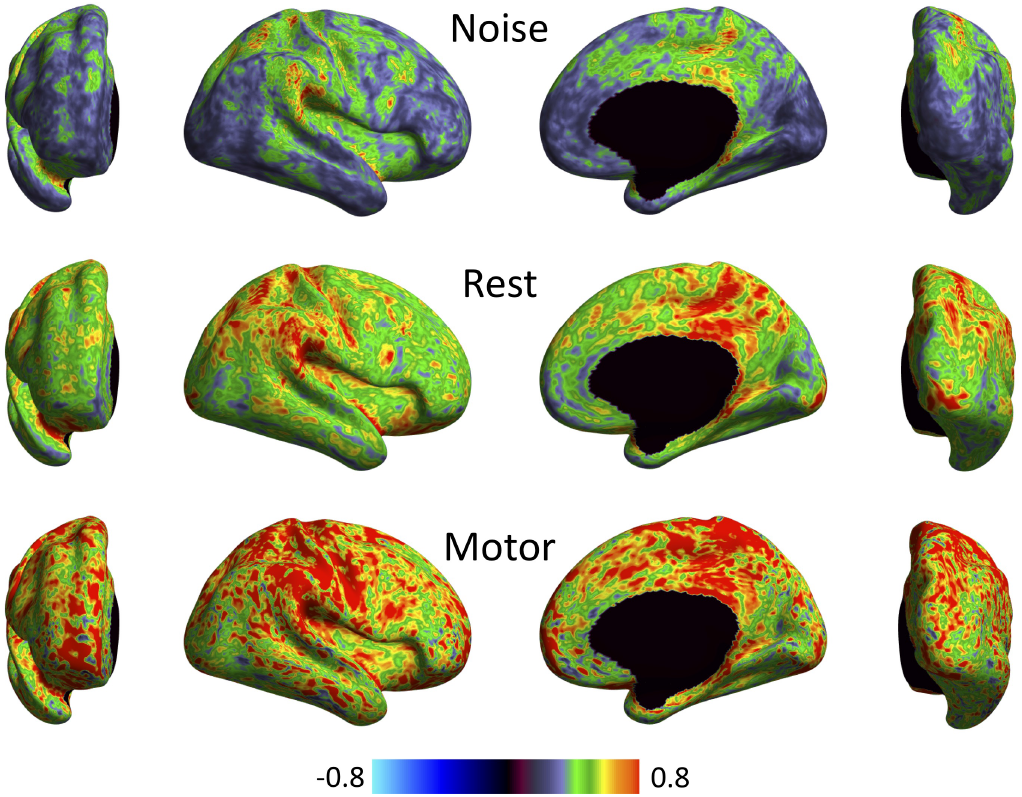
A delineation of the observed anomaly. The presence of local correlations that mirror the pattern of gyral folds emerges clearly, even in the case of noise. The figure shows surface-based, vertex-wise neighborhood correlation maps for three separate sets of volumetric data: noise (top row), resting-state (middle) and task data (bottom). Volume-to-surface mapping was carried out using cubic interpolation and the direct approach

A related phenomenon was first noted in (Glasser et al, 2013). The authors observed that increasing the resolution of functional MRI data decreases “geometric effects” on surface data. Their primary concern was, however, the implications this had for the correlation between relatively (geodesically) distant cortical areas.

In order to explore the phenomenon in the context of neighborhood correlations more rigorously, we used uncorrelated noise (drawn from a standard normal distribution) to create synthetic fMRI timeseries at the voxel level. These were then projected onto the surface using various volume-to-surface mapping approaches.

A measure of local connectivity was obtained by calculating the pairwise Pearson correlation between any two vertices within a local region (where a *local region* is defined as consisting of a given vertex and its neighborhood i.e. any vertices directly adjacent to it). Next, we performed the Fisher-z transform, and adopted the mean of z-scores as the local connectivity index for the given vertex.

The presence of regional boundaries that follow gyral patterns emerged when using both real and stochastic data. This prompted a more systematic exploration of the issue, with the aim of understanding and mitigating the problem.

## 3 Methodology

T1-weighted data from the minimally preprocessed Human Connectome Project Young Adult data set were used for this study (Glasser et al, 2013). These data were selected for the high quality of their acquisition, as well as to ensure state-of-the-art preprocessing (up to the point where the discussed issue is encountered); we considered a single subject. Noise timeseries were generated at three isotropic resolutions (0.7 mm, 1.4 mm and 2 mm) in volumetric space and pushed to the surface using five different volume-to-surface algorithms provided with Connectome Workbench (Marcus et al, 2011). The first two use simple trilinear and cubic interpolation at the level of the midthickness surface. The second two employ the advanced ‘ribbon-constrained’ technique, which makes use of constructed polyhedra constrained by the pial and white matter surfaces (one polyhedron per vertex). The amount of overlap between a given vertex’s polyhedron and any nearby voxels serves to weight the contribution of these voxels in the sampling process (in the “thin columns” (TC) variant, the polyhedra of neighbouring vertices do not overlap). The final approach, termed “enclosing”, simply uses the voxel directly beneath each vertex of the midthickness surface. In order to isolate the problem, and thus ensure it really arises from the interpolation of voxel data for surface mapping, random noise was generated independently for the vertices in the reconstructed surface; we refer to the corresponding model as the “null model”.

To guarantee a rigorous evaluation of the effect of all the different volume-to-surface mapping approaches, we carried out the following:

1. Volume-to-surface mapping of noise timeseries onto the native high-resolution surface (in the MNI standard space as per HCP data), using the five methods detailed above. Then down-sampling the maps to the surface resolution of interest (32k and 10k). We term this approach the “traditional approach”;
2. Volume-to-surface mapping directly onto surfaces having the required resolution (native, 32k and 10k), using the five approaches. We refer to this as the “direct approach”;
3. Volume-to-surface mapping onto midthickness surface meshes that approximate the resolutions of interest, but that have been modified to reduce the variance in inter-vertex distance (Attene, 2010). In this case, we employ the three most basic methods (trilinear, cubic and enclosing);
4. Construction of scatter plots illustrating the relationship between mean distance of neighborhood vertices and local correlations (refer to Figure 2). We also plotted histograms of the local correlations for all the different approaches (Figure 3).

**Fig. 2.**
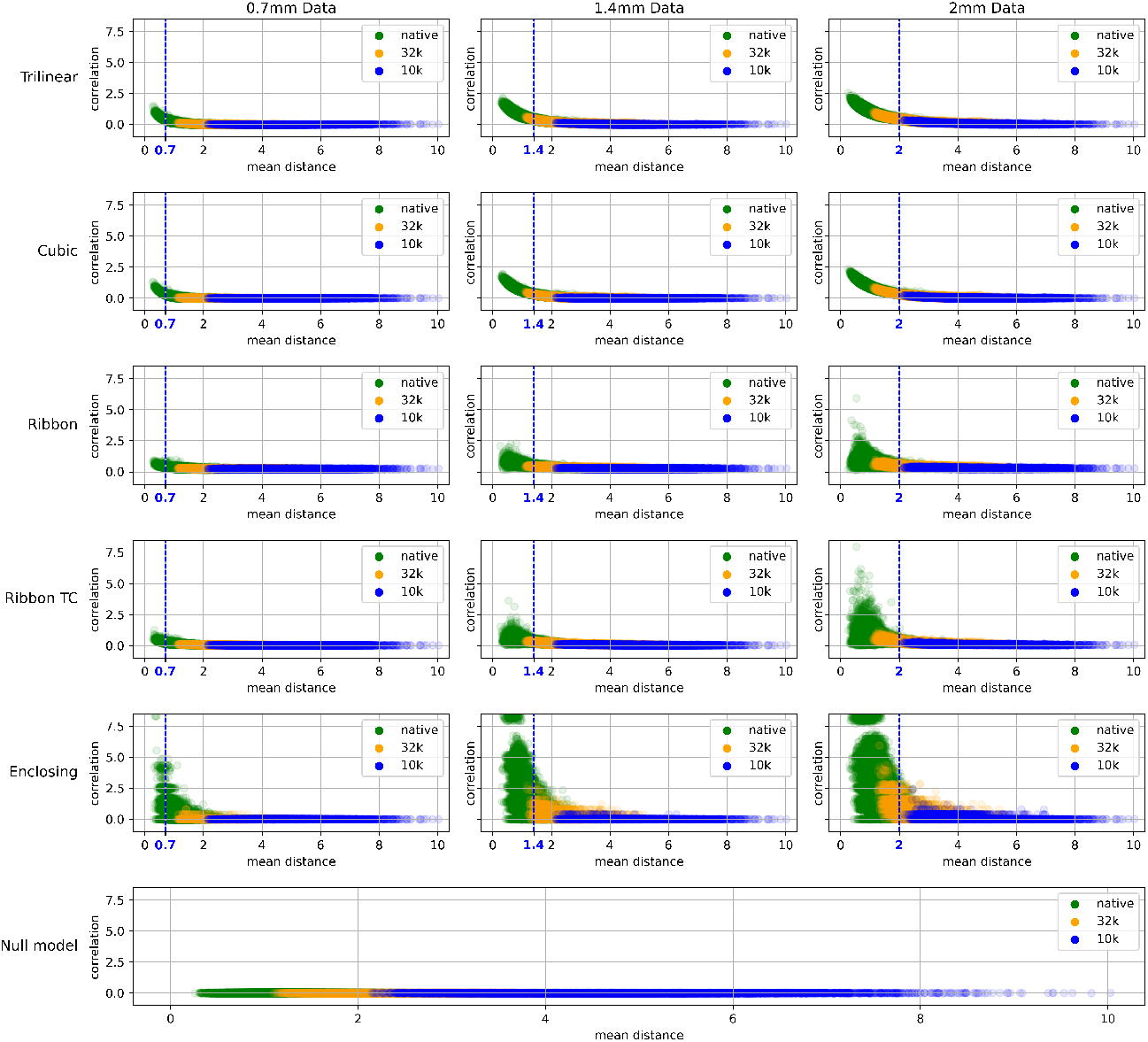
Variation of local correlation with mean intra-vertex distance for various direct mapping approaches and different underlying voxel resolutions (stochastic input in all cases). Data from medial wall structures were not included. The bottom graph shows what is expected if local correlations are computed for noise timeseries generated directly on a surface mesh (with no volume-to-surface mapping artefacts). The vertical dashed (blue) lines indicate the resolution of the underlying volumetric data in one dimension (the voxels are isotropic). As the mean distance between vertices approaches this resolution, geometric effects cause the average local correlation to increase

**Fig. 3.**
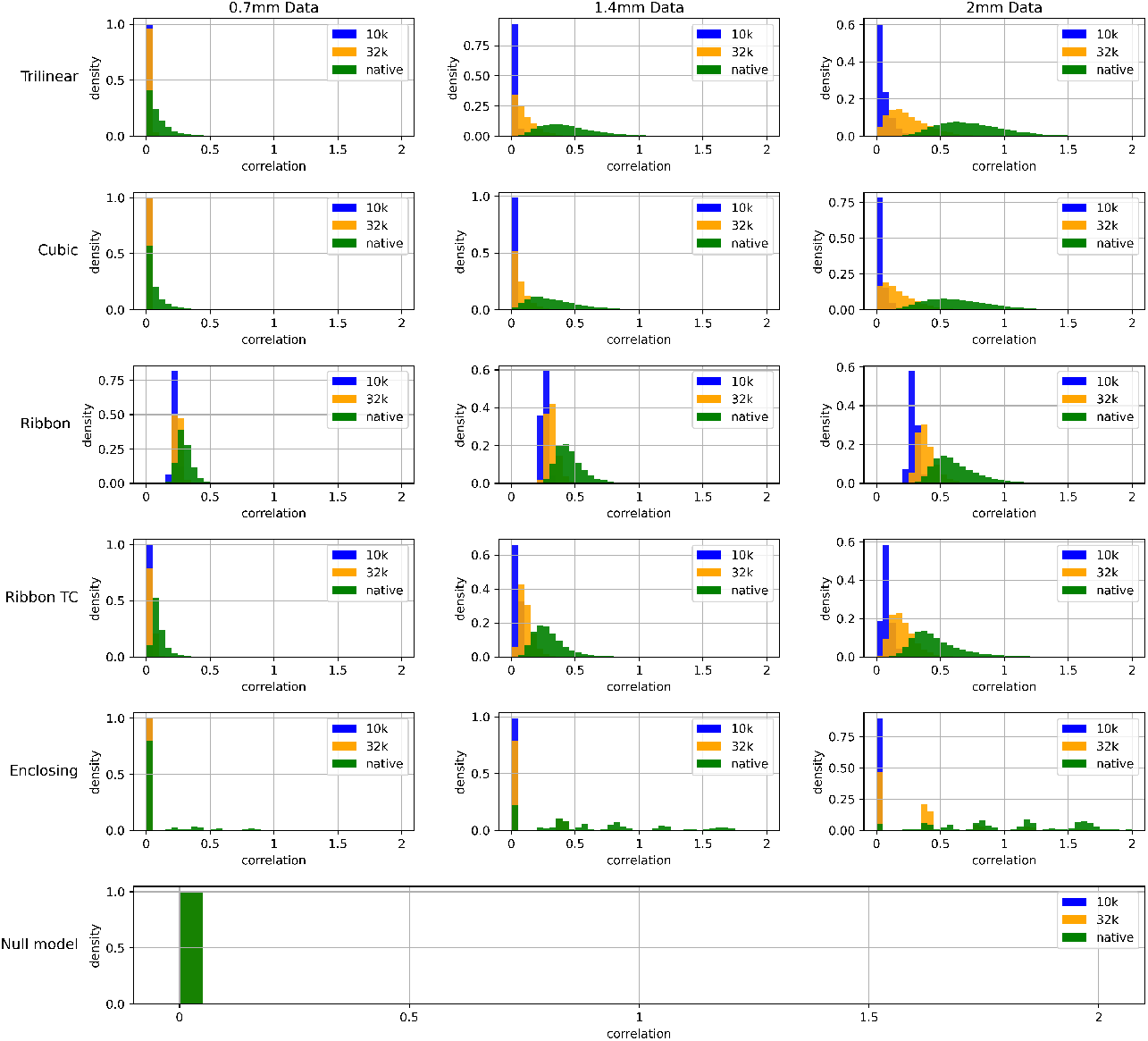
Variation of vertex probability density with surface-based local correlations computed from stochastic volumetric input data. We considered various direct mapping approaches and voxel resolutions. Midline structures were not taken into account for the purpose of producing these histograms

For the sake of brevity, only the most pertinent results are discussed in the following section. However, all results (including those for the traditional approach) and code are made available for further investigation and scrutiny and may be found at https://osf.io/dxr6f/.

## 4 Results

A clear relationship between surface resolution and the size of underlying volumetric elements (voxels) was found. As surface resolution (vertex density) is increased, or voxel data resolution decreased, more surface vertices sample the same voxels from the volume. In both the traditional and direct approaches, this results in an artefactual local correlation despite the original, volumetric data being uncorrelated noise. We note that this shortcoming should not be unique to brain data or surface-based processing; upsampling any data should present a similar issue. Figure 2 shows, for the direct approach, that local correlations are approximately zero until the mean distance between neighboring vertices decreases below a certain range (approximately equal to the resolution of the volumetric data).

## 5 Mitigation

From the results of our investigations two points emerge (in the context of local correlation analysis):

1. the resolution of the underlying voxel data should be higher than that of the surface mesh, or at least comparable to it. The important point is that the underlying data should not be of lower resolution than the surface mesh;
2. any variance in intra-vertex distance on the mesh will create systematic “problematic areas”. As a result, the uniformisation of surface meshes may mitigate the problem.

We ran the same pipeline on uniform meshes. As can be deduced from Figures 4 and 5, projecting data at a typical high fMRI resolution (2 mm) onto a 10k *uniform* surface greatly improves the local connectivity estimation of the data. Figure 5 (top) shows that the noise surface-to-volume mapping does not give rise to any obvious geometric patterns in local correlations.^1^ Further, additional analysis using real resting-state or task-based fMRI data reveals high local correlations in areas that conform to the expected regions of activity. It is important to note, however, that this approach comes at the cost of 1) decreasing spatial resolution and 2) adding extra steps to the pipeline (needed to regain vertex correspondence to a standard template).

**Fig. 4.**
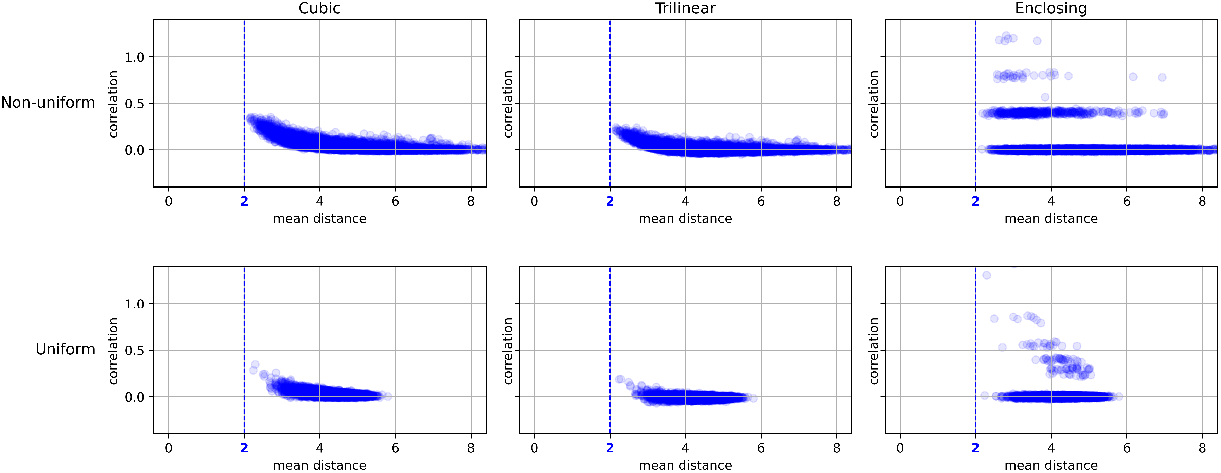
The impact of uniformisation of the mesh. Neighborhood correlations remain around zero (except for a small proportion of neighborhoods sampled with the enclosing approach) for all three volume-to-surface methods. In each case, we used stochastic 2 mm data, and mapped them to a surface with a resolution of 10k vertices

**Fig. 5.**
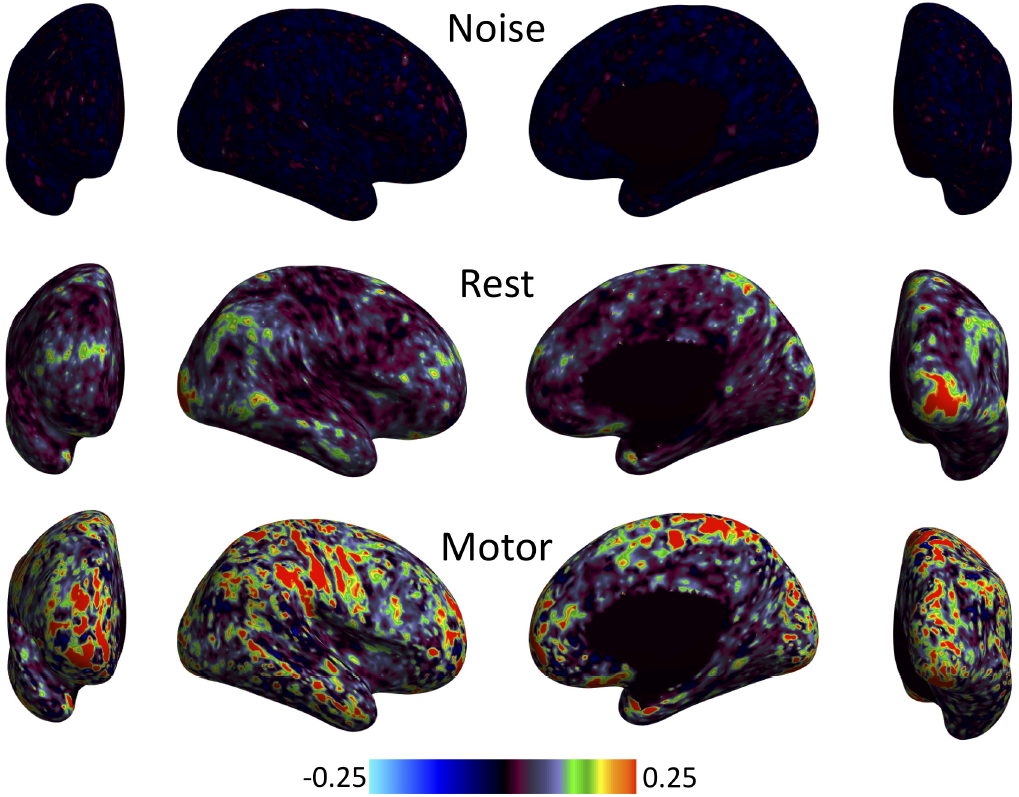
These are the final neighborhood correlation maps, obtained using the direct approach (cubic interpolation) on uniform 10k meshes. The difference between Figures 1 and 5 emerges clearly upon comparison

## 6 Conclusion

In this work, we note that:

1. the difference between the cortical surface vertex density and underlying voxel resolution is important to take into account;
2. not considering the implications of oversampling when mapping volumetric data to a relatively dense mesh has the potential to affect studies that deal with local neighborhood connectivity;
3. the artefactual geometric patterns correlated with the gyral structure of the cortex can be mitigated (but not necessarily eliminated) by a) mapping data to a lower resolution mesh and b) ensuring that the intra-vertex distance of the mesh is of relatively low variance.

In summary, this study highlights the occurrence of geometric effects in surface-based analyses of local neighborhood connectivity. We caution that any researcher interested in investigating local correlations in brain function should take such effects seriously.

## Acknowledgments

The authors would like to acknowledge Lucas Q. Costa Campos for fruitful discussions on multiple versions of this manuscript. We would also like to thank Alexander Opitz and Michael Milham for useful suggestions during the initial investigation.

## Declarations

### Ethics approval

This research satisfied the University of Malta’s research ethics committee (UREC) self-assessment criteria.

### Availability of data and materials

Data were provided by the Human Connectome Project, WU-Minn Consortium (Principal Investigators: David Van Essen and Kamil Ugurbil; 1U54MH091657) funded by the 16 NIH Institutes and Centers that support the NIH Blueprint for Neuroscience Research; and by the McDonnell Center for Systems Neuroscience at Washington University.

We made use of HCP Young Adult data (the S1200 release; see https://www.humanconnectome.org/study/hcp-young-adult/document/1200-subjects-data-release), processed by the HCP with the following pipelines:

– Original MSM-Sulc based preprocessing (v3.4.0)
– Spin Echo Bias Field Prerequisite Files (v3.12.0)
– MSM-All DeDrifting and Resampling based on MSM-All registration (v3.13.2)

Details about the subset of subjects the work is based on and further information about processing/analysis may be obtained from the authors.

### Code availability

The code required to recreate the complete analysis is available at https://osf.io/dxr6f/. We also provide a number of additional figures.

## Footnotes

1 We verified this by repeating the procedure for 23 other participants included in the data release. In all cases, similar results were obtained (we have made these available as supplementary material).

